# Design and Analysis of a Whole Body Non-Contact Electromagnetic Stimulation Device with Field Modulation

**DOI:** 10.1101/416065

**Authors:** Sergey N. Makarov, Gene Bogdanov, Gregory M. Noetscher, William Appleyard, Reinhold Ludwig, Juho Joutsa, Zhi-De Deng

## Abstract

This study describes a whole-body, non-contact electromagnetic stimulation device based on the concept of a conventional MRI Radio Frequency (RF) resonating coil, but at a much lower resonant frequency (100–150 kHz), with a field modulation option (0.5–100 Hz) and with an input power of up to 3 kW. Its unique features include a high electric field level within the biological tissue due to the resonance effect and a low power dissipation level, or a low Specific Absorption Rate (SAR), in the body itself. Because of its large resonator volume together with non-contact coupling, the subject may be located anywhere within the coil over a longer period at moderate and safe electric field levels. The electric field effect does not depend on body position within the resonator. However, field penetration is deep anywhere within the body, including the extremities where muscles, bones, and peripheral tissues are mostly affected. A potential clinical application of this device is treatment of chronic pain. Substantial attention is paid to device safety; this includes both AC power safety and exposure of human subjects to electromagnetic fields. In the former case, we employ inductive coupling which eliminates a direct current path from AC power to the coil. Our design enhances overall device safety at any power level, even when operated under higher-power conditions. Human exposure to electromagnetic fields within the coil is evaluated by performing modeling with two independent numerical methods and with an anatomically realistic multi-tissue human phantom. We show that SAR levels within the body correspond to International Electrotechnical Commission (IEC) safety standards when the input power level of the amplifier driver does not exceed 3 kW. We also show that electric field levels generally comply with International Commission on Non-Ionizing Radiation Protection safety standards if the input power level does not exceed 1.5 kW.

## I. Introduction

In this present study, we have designed and constructed a whole-body non-contact, electro-magnetics stimulation device based on the concept of an MRI Radio Frequency (RF) resonating coil. However, this has been accomplished at a much lower resonant frequency (100–150 kHz) and with a field modulation option (0.5–100 Hz) at input powers up to 3 kW. Due to a large resonator volume and its non-contact nature, the subject may be conveniently located anywhere within the resonating coil over a longer period of time at moderate and safe electric field levels. The electric field effect does not depend on a particular body position within the resonator. The field penetration is deep everywhere within the body including the extremities; muscles, bones, and peripheral tissues are mostly affected. Over a shorter period of time, the electric field levels may be increased to relatively large values with an amplitude of about 1 V/cm.

The primary potential clinical application for the device is treatment of chronic pain. Approximately 20% of the adult population report chronic pain and approximately one fifth of them have neuropathic pain characteristics [1–4]. Common forms of chronic pain include nociceptive pain, which involves stimulation of pain receptors secondary to tissue damage, or neuropathic pain, which involves injury or dysfunction of the central or peripheral nervous system [5]. Neuropathic pain is often rated as particularly intense and distressing and can have a significant negative impact on activities of daily living and quality of life [1–4]. A commonly prescribed treatment for chronic pain is opioids [6]. However, the use of opioids for pain relief is controversial owing to concerns about addiction and misuse [7].

One known yet not very efficient electromagnetic treatment of neuropathic and other pain conditions is Transcutaneous Electrical Nerve Stimulation, or TENS. It typically relies upon a portable unit which applies low-level electrical currents through electrodes attached to the skin. However, proper placement of electrodes and mitigating skin irritation are two major points of concern [4]. A well-known problem with electrodes attached to the skin is a strong regional current injection close to the contact area but not necessarily close to the deeper treatment region and may result in burning effects in the vicinity of electrodes where deep penetration is required. A number of systematic reviews of the effect of TENS on various painful conditions such as neuropathic pain, phantom limb pain, fibromyalgia, labor pain, rheumatoid arthritis, chronic lower back pain, etc. are available [1, 4, 8–12].

An alternative to electrical stimulation with contact electrodes is magnetic stimulation, which induces electric field in the body via electromagnetic induction thus alleviating the issue of current shunting in the skin. The major goal of this paper is to introduce a distinctive, noncontact yet potentially powerful full-body electromagnetic stimulation device concept, including

i. the underlying physical model in the form of an electromagnetic resonator;
ii. the hardware design, preliminary measurements, and tests;
iii. a detailed computational analysis of the electric fields induced in the body along with safety estimates.

The key idea of our stimulator is the creation of an initial rotating magnetic field that is routinely produced in resonant MRI RF coils, but for a completely different purpose, namely atomic spin excitation and RF signal acquisition. Furthermore, the resonant frequencies used in MRI RF coils are very high, typically 64 MHz or higher, whereas our coil operates at much lower resonant frequencies (100–150 kHz). Nonetheless, an RF birdcage coil [13, 14] seems to be an ideal starting point for our design. When the frequency becomes low as in the present case, the standard RF birdcage coil will possess very low inductance. Tuning such a birdcage toward resonance at low frequencies would require large capacitances. This, however, means a low *Q*-factor (quality factor *Q* is the “gain” of the resonator) and higher costs, as well as higher fabrication complexity [15, 16], and will restrict the use of the conventional birdcage coil to frequencies above at least a few megahertz. Different methods to overcome this difficulty have been suggested [15–20], but they are all limited to small-size coils.

Our design utilizes a unique large-scale low-pass birdcage coil architecture with a large number (144) of long rungs and bridging capacitors, and possesses a superior quality factor of approximately 300. The design does not use any magnetic materials; it is robust, lightweight, and portable. However, it achieves quite significant electric field levels within the body due to the resonance effect. This allows us to use relatively low input power levels and standard low-cost power electronic equipment. Although the resonant or carrier frequency is fixed, the electric field within the resonator volume can be amplitude modulated from approximately 0.5 Hz to 100 Hz.

The study is organized as follows. Section II describes a theoretical device model and specifies field distribution within the resonator. Section III describes hardware design, test, and functionality, including semiautomatic operation/tuning and representative continuous run times. Section IV provides computational results for the electric field distribution within the body obtained via two independent numerical methods. Section V discusses possible application scenarios as well as device modifications. Section VI concludes the paper.

## II. Theoretical model of the stimulator

### A. Concept of the magnetic stimulator. Two-dimensional analytical solution

Fig. 1 shows the anticipated device concept. An external uniform rotating (or circularly-polarized) magnetic flux density **B** with amplitude *B*_0_ is created in the transverse plane around a tissue volume, depicted in Fig. 1a. By Faraday’s law of induction, this field excites a axial, rotating electric field **E** in free space or in a tissue volume, which is expressed in terms of the magnetic vector potential **A**,

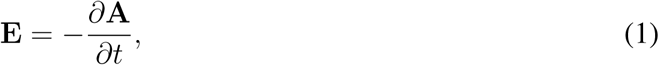

as shown in Fig. 1b; we set **B** = ∇×**A**. Thus, when a biological body is placed into this volume, a significant non-invasively excited electric field in the *axial* direction will appear parallel to the major peripheral nerves, spinal cord, long bones, major arteries, veins and other structures. This is in contrast to a solenoidal coil wound around the body creating electric fields and currents in the less desirable transverse plane.

**Fig. 1.**
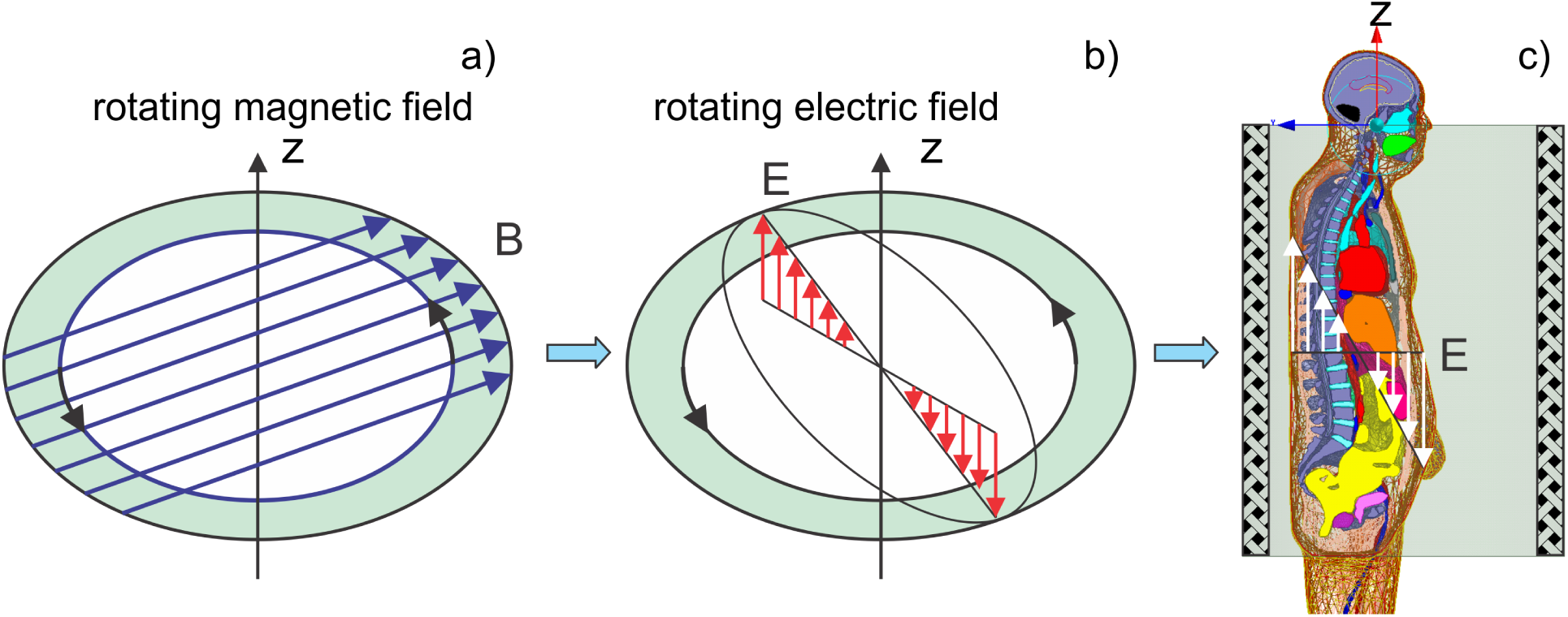
Concept of non-invasive electric field excitation via the induction mechanism. a) A rotating magnetic field shown in excites the b) rotating electric field within the conductor and c) within the body.

The rotational character of the field also assures that not only one body cross-section (e.g. coronal or sagittal) will be subject to the electric field excitation, but the entire body volume.

In the ideal, two-dimensional case and for any conducting target with a strict cylindrical symmetry placed into the device, either homogeneous or not, the corresponding two-dimensional problem, shown in Fig. 1a,b will have an exact analytical solution in the quasi-static (or eddy current) approximation. The electric field within the target is given by [21]

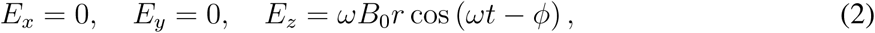

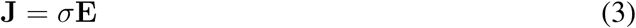

where *r* is the radial distance from the coil axis in cylindrical coordinates, *ω* is the angular frequency, *ϕ* is an arbitrary phase, **J** is induced current density, and *σ* is the (local) medium conductivity, which is either constant or obeys cylindrical symmetry. Although Eqs. (2)–(3) might be used to roughly estimate the electric field in the body based on a cylindrically symmetric assumption, its actual value will deviate as shown below.

### B. Three-dimensional coil model

The external rotating magnetic flux density **B** is created using a volumetric resonator in the form of a low-pass birdcage coil. This design is well-known in the magnetic resonance radio-frequency coil community [13, 14]. We modified the design to operate at a much lower frequency of 100 kHz and with a very high loaded quality factor.

A computational model of a particular resonator constructed in this study is shown in Fig. 2a with the electric current distribution to scale. The coil has a diameter of 0.94 m and a length of 1.10 m; the coil resonates at 100 kHz or at 145 kHz depending on the values of the bridging capacitors. The coil consists of two rings (top and bottom) joined via multiple straight rungs, each bridged with a lumped capacitor at its center. The capacitors control the coil’s resonant frequency. The resonating coil is fed via two lumped ports in quadrature, or using inductive coupling with two loops in quadrature as explained below.

**Fig. 2.**
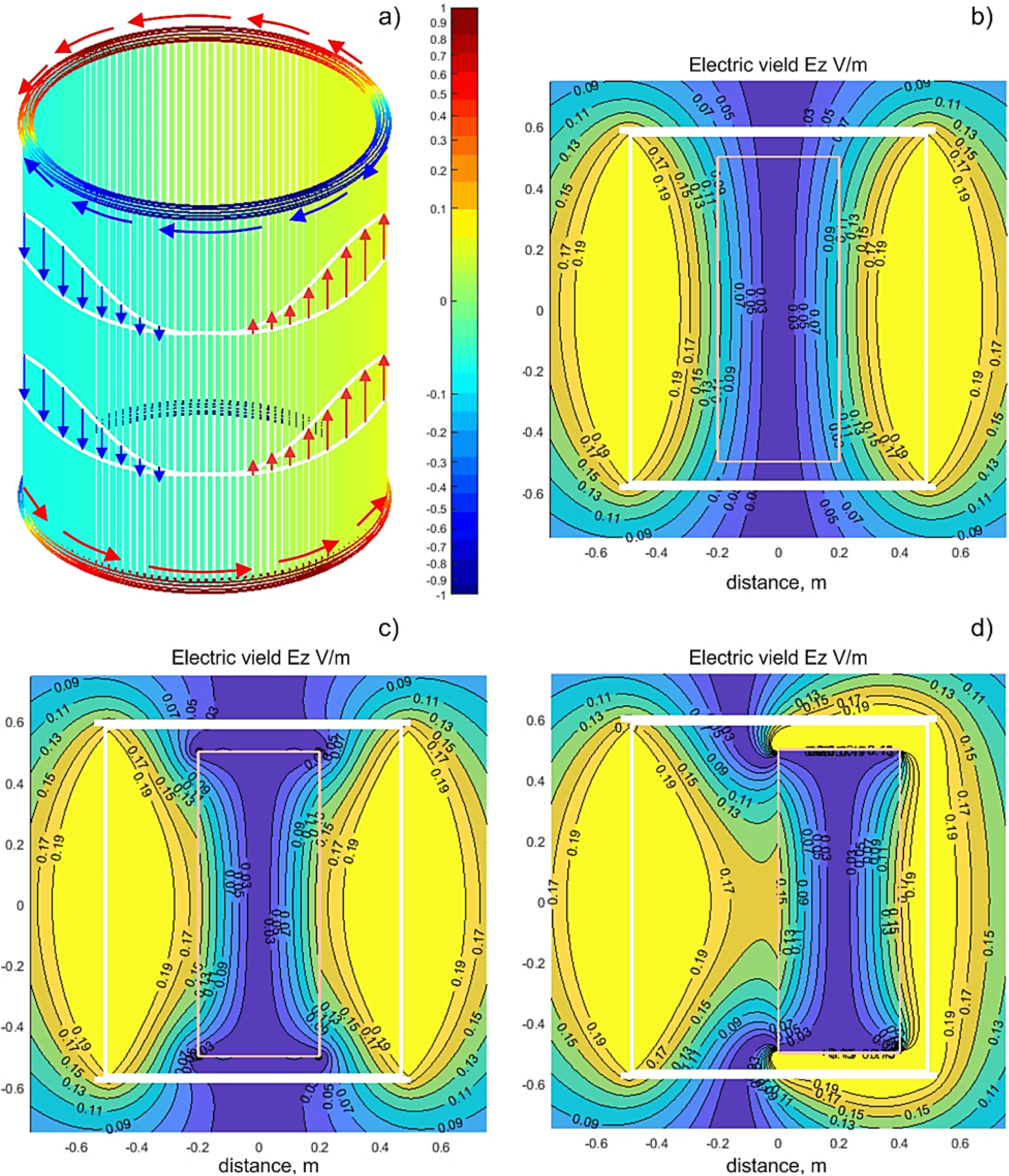
Current distribution in the coil resonator and the associated electric field created by the coil when the ring current amplitude is 1 A for one resonant mode. a) Electric current distribution in the coil along with the current colorbar to scale; b) magnitude of the vertical electric *E*_*z*_ in V/m for an empty coil in the coronal plane; c) magnitude of the vertical electric *E*_*z*_ in V/m for the coil with a coaxial conducting cylinder 0.4 *×* 1 m inside; c) magnitude of the vertical electric *E*_*z*_ in V/m for the coil with a conducting cylinder 0.4 *×* 1 m shifted in the transverse plane inside.

From the modeling point of view, the resonant electric current in both rings at any fixed time instant behaves like a full period of a sine function of polar angle *φ*. This ring current distribution is shown in Fig. 2a. As time progresses, the ring current distribution shown in Fig. 2b rotates with angular frequency *ω*. As a result, the time-domain ring current *i*(*t, φ*) in the top and bottom rings can be expressed in the form

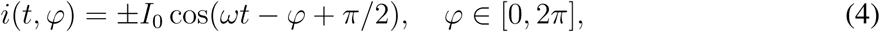

where *I*_0_ is the current amplitude determined by the excitation power and by the quality factor (or the “gain”) of the resonator.

The AC current in each rung shown in Fig. 2a does not change along its length. Simultaneously, at any fixed time instant, it also varies from rung to rung as a harmonic function of the polar angle *φ* with the full period corresponding to the ring circumference. This rung current distribution is shown in Fig. 2a. As time progresses, the rung current distribution shown in Fig. 2a also rotates with angular frequency *ω*. Each individual time-domain rung current density *j*(*t, φ*) can be expressed in the form

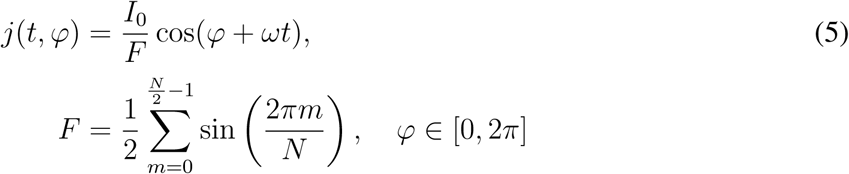

where *N* is the total number of rungs. This form obeys the current conservation law, or Kirchhoff’s Current Law (KCL), at every ring–rung junction.

The useful current, which creates a nearly constant horizontal rotating magnetic field *B*_r_ with amplitude *B*_0_ and axial rotating electric field *E*_*z*_ according to Eq. (2), is the rung current density *j*(*t, φ*). Contributions of each rung add up in a constructive manner. The ring current, on the other hand, does not contribute to the axial (or vertical) electric field, *E*_*z*_. However, it may create a strong transverse electric field very close to the rings.

It should be pointed out that Eqs. (4) and (5) describe the rotating current behavior, which is a combination of two elementary resonant modes. Each elementary mode does not rotate and appears as depicted in Fig. 2a. However, when excited in quadrature (with a 90 degree phase shift and a 90 degree excitation offset along the coil circumference), both modes combine to create the current distribution given by Eqs. (4) and (5) and the associated rotating electric field. The rotation phenomenon enable us to treat the entire body and not merely a singular component or region.

### C. Electric field model with and without a simple conducting object

Fig. 2b–d show the resulting electric field distribution in the coil (coronal plane) when the current amplitude *I*_0_ = 1 ampere in either ring given by Eq.(4). The results are given for one resonant mode is shown in Fig. 2a. Due to linearity, this result can simply be scaled for other excitation levels. Accurate field computations have been performed with the fast multipole method described in [22]. The magnitude of the axial component *E*_*z*_ in V/m for an empty coil is shown in Fig. 2b. The electric field is indeed zero at the coil’s center.

When a conducting object representing a load is inserted into the coil, the field distribution changes. Fig. 2c shows the distribution when a conducting cylinder with a diameter of 0.4 m and a length of 1 m is inserted into the coil along its axis. The particular conductivity value *σ* does not matter since only the conductivity contrast, (*σ − σ*_air_)*/*(*σ* + *σ*_air_), is present in the solution [22]. This value is always unity since *σ*_air_ = 0.

An interesting and useful effect is observed in Fig. 2c: we see a “pulling” of the electric field into the cylinder close to the coil center. This is due to surface charges that appear at and near the cylinder tips. As a result, the electric field close to the cylinder surface at the center plane of the coil increases by nearly 36%.

Another remarkable observation (this effect is common in MRI RF coils) follows from Fig. 2d where the conducting cylinder has been shifted from the coil axis to the right by 0.2 m. While the electric field outside the conducting cylinder clearly changes, the field within the cylinder remains nearly the same, as observed in Fig. 2c. This may be explained as a result of the electric field being induced by the magnetic field, similar to eddy currents. Since the magnetic field is relatively homogeneous in the transverse plane of the coil, the induced electric fields in a load should not strongly depend on the transverse position of the load within the coil.

These rudimentary simulations allow us to establish two basic facts relevant for the analysis of realistic electric field distributions in a human body within the coil. First, we expect that the average transcutaneous electric field will be slightly higher than predicted by the air-filled coil model in dorsal, abdominal, and lumbar body regions. Second, we expect that the field within the body will not change significantly when the body is moved within the coil in the transverse plane; this circumstance seems to be quite useful from a practical point of view.

Obviously, an accurate field evaluation with a realistic human model is desirable; we will present our modeling results in Section IV.

## III. Hardware design and test

### A. Power amplifier/driver

In order to create the rotating magnetic and electric fields as seen in Fig. 1a–b, two resonant modes are excited in the coil resonator. These modes display the same current distribution as shown in Fig. 2a, but rotated by 90 degrees about the coil axis with respect to each other as well as having an additional temporal phase shift of 90 degrees.

To accomplish this, a custom designed class-D, high-efficiency, single-frequency power amplifier (PA), whose circuit schematic is presented in Fig. 3a, was constructed and prototyped as shown in Fig. 3b–c. The upper block in Fig. 3a is a class-S modulator, which is followed by two class-D output stages in quadrature exciting the two resonant modes. The PA has two outputs for each resonant mode and generates a harmonic power RF signal at a fixed carrier frequency at each output. This frequency is typically around 100 kHz. At the same time, the PA may be tuned to operate at any carrier frequency from 30 kHz to 300 kHz in the LF band. The PA operation, including variable power levels, an optional variable modulation frequency, and a semiautomatic patient-specific RF frequency tuning procedure, which is automated via a microcontroller board, can be seen in Fig. 3b.

**Fig. 3.**
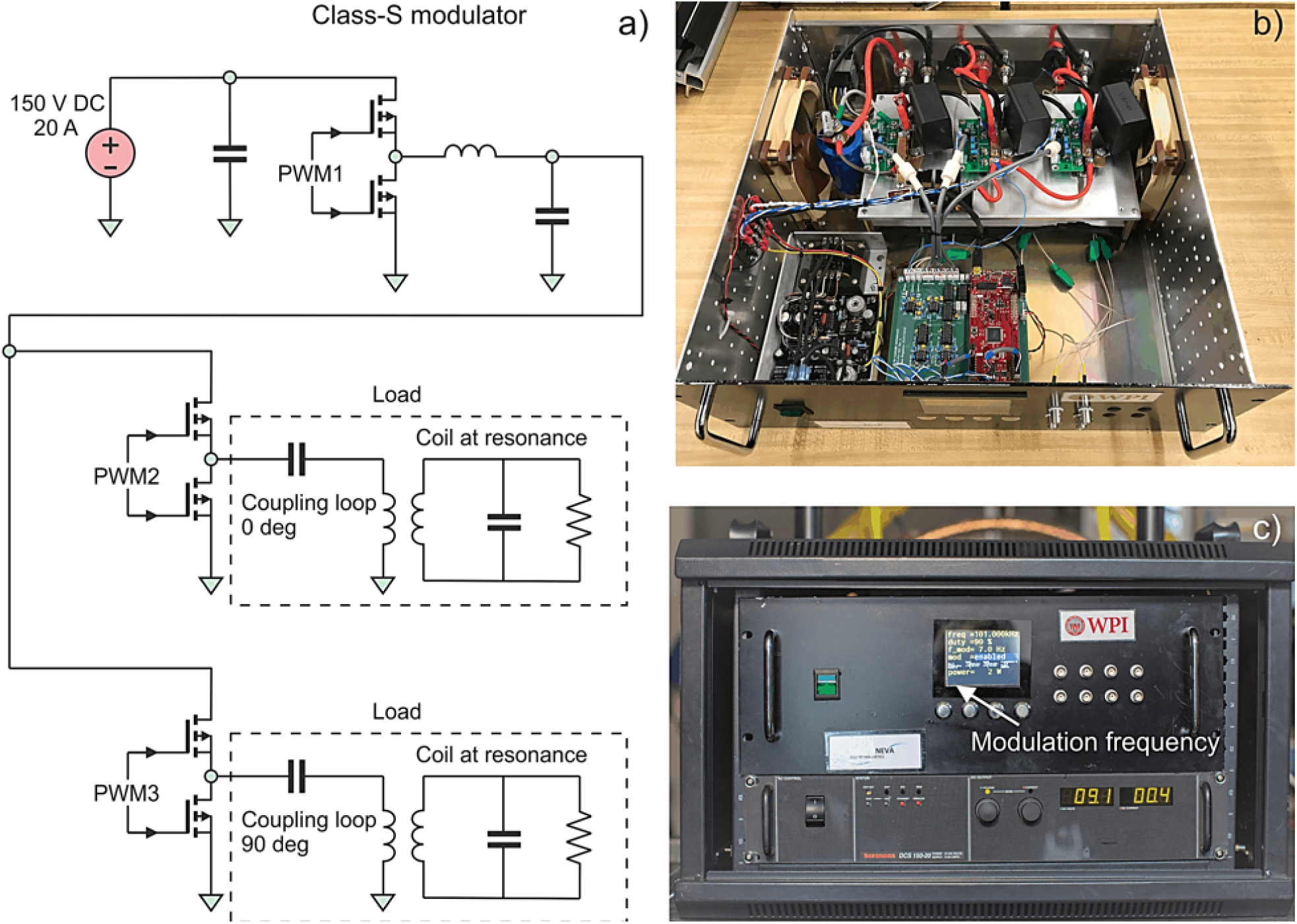
a) High-level circuit schematic of the two-channel power amplifier; b) rackmount air-cooled assembly of the electronic hardware; c) amplifier display controlling output power and modulation frequency (if used).

The PA output stage is powered by a 3 kW Sorensen DCS 150-20 Variable Regulated DC (Direct Current) power supply seen at the bottom of Fig. 3c. When connected to a standard three-phase 208 VAC outlet, the max RF output power is about 2.9 kW, based on 3 kW DC power. Alternatively, when connected into a single-phase 240 VAC outlet, the max RF power reduces to about 2.3 kW based on 2.4 kW DC power.

Arbitrary *modulation* (pulse or CW) of the carrier signal with a maximum modulation frequency component of 1 kHz is available via the modulator. The modulation bandwidth is limited mainly by the coil envelope time constant of about 1 ms. Typical modulation is sinusoidal in the 0.5 Hz to 100 Hz range, generated by the PA firmware.

The PA also monitors its output power and load impedance. It uses this information to automatically adjust the carrier frequency in a narrow band such that the output power remains on target. The amplifier cost including the DC power supply is under $10,000. The prototype 100 kHz PA was assembled in a rackmount case shown in Fig. 3b–c.

The reason for designing a custom, fixed-frequency PA is the lack of an affordable and appropriate commercial model. Industrial low frequency RF power supplies, e.g., Comdel’s CLB3000, are costly and require a matched 50Ω load. Because our load impedance varies widely with frequency, keeping the load matched is a challenge. It would require load impedance monitoring and fine frequency control (potentially difficult with a commercial unit), and/or a software-controlled matching network (costly). Additionally, generating two outputs in quadrature would require either a 90° hybrid (another costly component), or phase-locking two commercial PAs at a 90° phase difference, which can be difficult. Finally, the majority of commercial PAs require water cooling, whereas our PA relies on air cooling. One disadvantage of our custom design is its unknown reliability, a factor that will be proven over time.

### B. Coupling and matching power amplifier to the resonating coil

The amplifier is coupled to the resonating coil inductively via two proximate loops. One such loop is shown in Fig. 4c. Apart from certain technical advantages of the inductive coupling, this methodology assures that there is no direct current path from the AC power outlet to the coil. This design enhances overall device safety at any power level, including high-power operation.

**Fig. 4.**
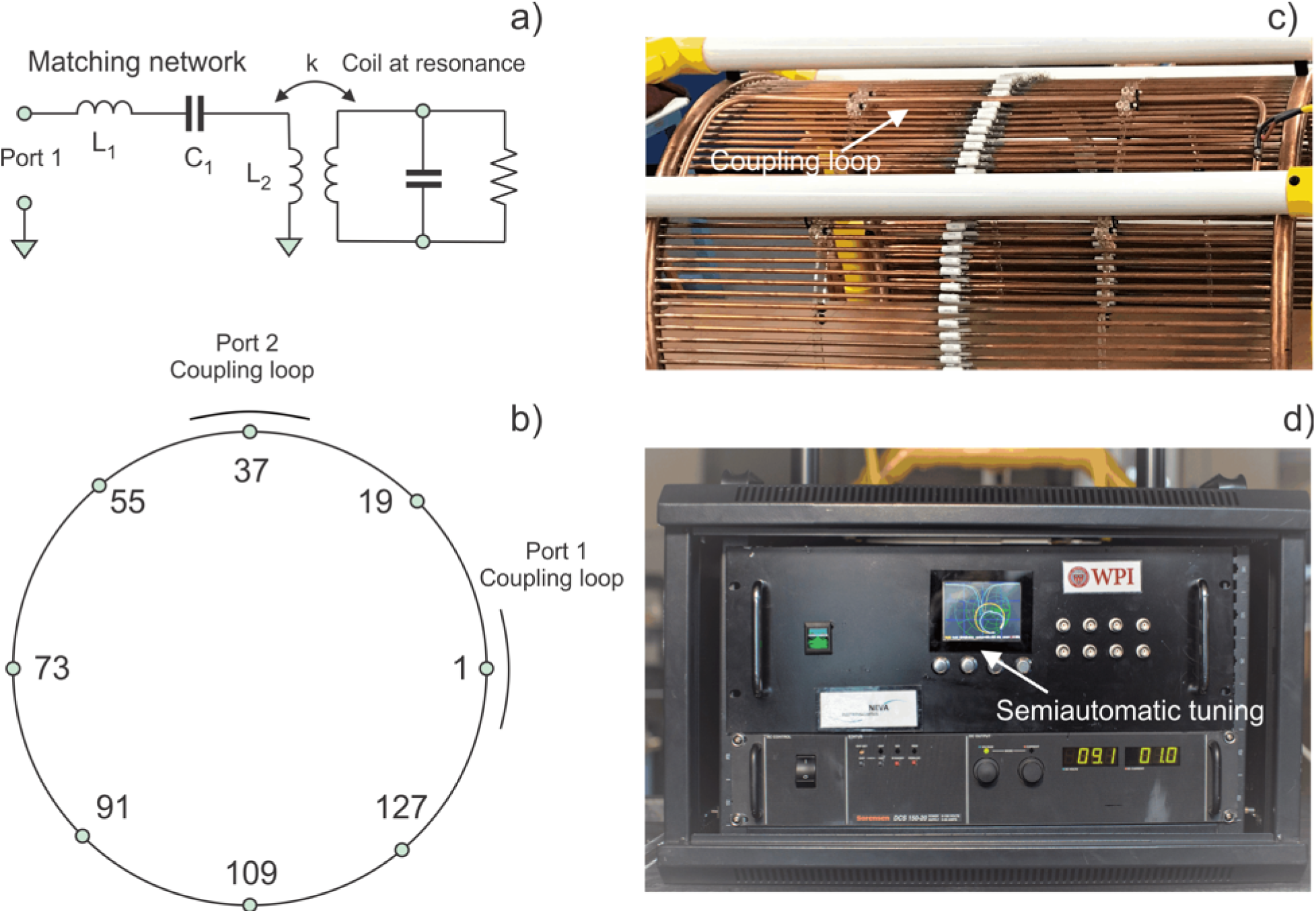
a) Matching and tuning network of the power amplifier; b) assembly of two coupling-loop feed around the coil circumference with 144 rungs; c) non-contact inductive coupling of the power amplifier to the coil resonator at one port; d) Smith chart/reflection coefficient display of the power amplifier controller used for semiautomatic tuning at any desired time instant.

The matching network for a single coil port is shown in Fig. 4a. Two ports with identical matching networks are located 90° apart around the coil structure, as shown in Fig. 4b. The port matching network consists of a series capacitance *C*_1_, series inductance *L*_1_, and the fixed inductance *L*_2_ of the inductive loop shown in Fig. 4c.

The matching network is tuned such that the load looks mostly resistive over a small frequency band around the coil’s resonance. For example, the load reactance stays quite low from 99.85 kHz to 100.15 kHz, while the resistance varies from 1.3Ω to 6Ω. Because the coil resonance shifts as the coil heats up, the operating frequency must be actively adjusted to compensate for this change, or the output power will vary.

Another important safety feature of the matching network is its benign behavior when subjected to a step response in output power. The matching network avoids large spikes in PA output current while energy is building up in the resonating coil.

Finally, the matching network presents a sufficiently inductive impedance to higher harmonics of the PA output voltage. This protects the output stage, and ensures that voltage transitions occur when the output current is low, thereby improving efficiency. The efficiency of the PA with the expected load is estimated to be greater than 90% over a wide output power range.

### C. Tuning procedure

The semiautomatic tuning procedure ensures that the reflection coefficient of both modes stays below −25 dB when matched to the maximum-power coil impedance of *Z*_0_ = 2.5Ω and that both resonances are within a 20 Hz band. The primary adjustable components are the series capacitance *C*_1_ in Fig. 4a and the coil rung capacitors at the numbered locations in Fig. 4b.

The tuning procedure is controlled and guided by the Smith chart/reflection coefficient display of the PA controller seen in Fig. 4d. It includes a number of well-defined steps, and is applied to the coil at its designated operating location in an effort to account for the presence of large nearby metal objects. The tuning procedure is simple to perform.

### D. Coil assembly, complete device setup, and operation

A resonator coil prototype made of thin-walled, light copper tubing was constructed; it is shown in Fig. 5a–b. Tubing thickness was kept at 0.75 mm or greater to avoid excessive eddy current losses in the copper. The capacitor size in Fig. 5a is relatively large since those components must operate at significant currents levels, up to 15 A RMS per rung for a maximum power of 3 kW, and at large voltages, up to 240 V RMS for a maximum power of 3 kW across the capacitor. The total coil weight without the frame is approximately 120 lbs (54 kg).

**Fig. 5.**
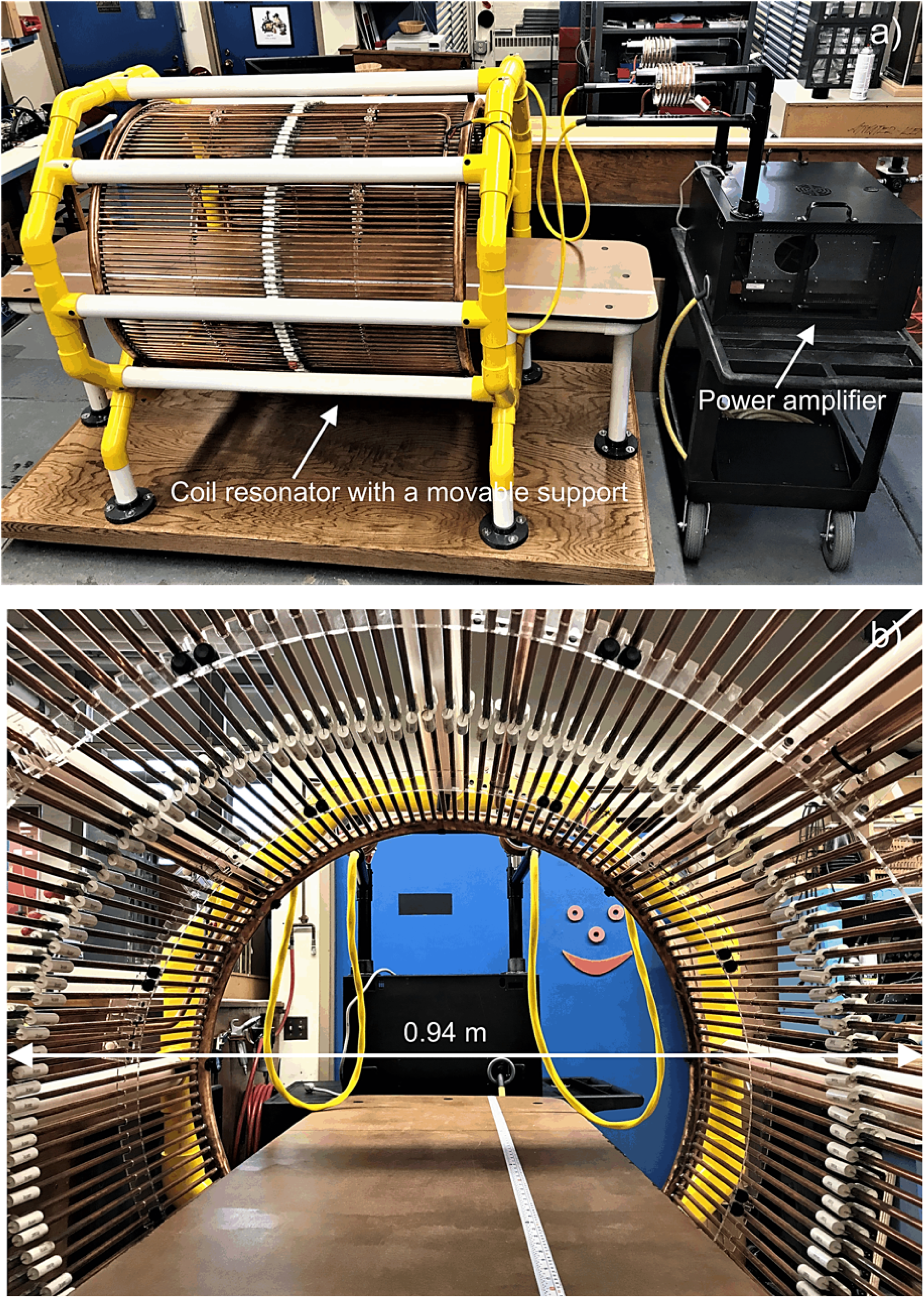
a) Complete framed and movable device setup; b) coil resonator assembly.

This durable coil prototype was then framed, augmented with a horizontal bed, and placed horizontally to enable a subject to rest in the coil as shown in Fig. 5a, which simultaneously shows the complete device setup. The entire coil frame is portable. The distance between the PA, which is connected to the inductive coupling loops of the coil via two isolated cables, can vary from 1 to 3 meters, although larger distances may be possible. As mentioned above, there is no direct ohmic current path from the AC power outlet to the coil which is an important safety feature.

An operator sets the power level, the modulation frequency, and performs RF tuning at the beginning of the resonator operation and for a particular coil load. Furthermore, an automatic RF tuning procedure has been implemented that takes coil heating into account. At the maximum power level, the ring conductors of the coil heat up to approximately 70–75°C at continuous operation.

Continuous coil operation at the maximum input power of 3 kW was tested multiple times with an uninterrupted operation time of up to 2 hours and with a cumulative operating time in excess of 100 hours.

### E. Quality factor of the resonator and the magnetic field strength

The achievable field strength in the coil is determined by three factors: the strength of the PA, the quality factor *Q* or the “gain” of the resonator and the coil volume. When the quality factor is high, large field values within the coil can be achieved at a modest input power.

When measured across one of its rung capacitors, the birdcage coil behaves like a parallel resonator in a narrow frequency range around the resonant mode. Using a setup with a signal generator and oscilloscope, the resonator’s quality factor has been estimated in the form:

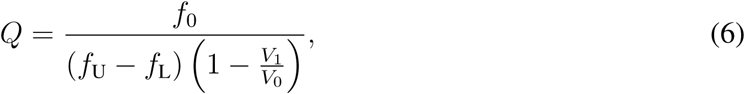

where *f*_0_ is the resonance frequency and voltage *V*_0_ is the open-circuited generator voltage. Voltage *V*_1_ is measured at resonance (where it is maximized), *f*_L_ and *f*_U_ are the lower and upper frequencies, respectively, where voltage *V*_1_ drops by 3 dB from its peak at resonance. This method is accurate in the high *Q* limit. The experimental data for 145 kHz is given in Table I. Table I reports a *Q*-factor value of about 300 at 145 kHz and a minimum difference between loaded (with a human body) and unloaded coil, which is to be expected at this low frequency. This value agrees with the theoretical/simulation prediction to within 10%. A similar quality factor (about 290) was obtained at 100 kHz.

**TABLE I.** Measured *Q*-factors for the coil resonator at 145 kHZ. Generator’s voltage is 1 V RMS. The load is a 200 lb. subject.

| Coil | $V_1$ (V RMS) | $f_0$ (kHz) | $f_L$ (kHz) | $f_U$ (kHz) | $Q$ |
| --- | --- | --- | --- | --- | --- |
| Unloaded | 0.6224 | 145.30 | 144.567 | 145.875 | 295.8 |
| Loaded | 0.6213 | 145.28 | 144.566 | 145.879 | 292.2 |

The established quality factor value is superior to the values reported in the literature for known low-frequency resonator coils (used for low-field MRI) in Table II. Note that all listed competitors have a much smaller coil size/volume and typically a lower quality factor.

**TABLE II.** Characteristics of existing low-frequency RF coils given for comparison with the present resonator prototype.

| Ref. # | Coil type | Frequency (kHz) | $Q$ (unloaded) |
| --- | --- | --- | --- |
| [15] | Wound birdcage coil (84 mm long and has a diameter of 73 mm) | 386 | 180 |
| [16] | Wound birdcage coils and a solenoid.<br>The diameter and the length of the coils are 70 mm | 238/425 | 100–280 |
| [17] | 27 turn saddle coil made of Litz wire with 8 cm diameter | 373 | 105 |
| [18] | 4-Coil Whiting–Lee configuration, 33 cm long | 83.6 | 100 |
| [19] | Solenoidal coils; 6–46 cm in length and 4–52 cm in diameter | 210/275 | 60–30,<br>reduced $Q$ |
| [20] | Cylindrical saddle-shaped loops (5 saddle pairs of 10 turns each), coil diameter is 26 mm | 87 | NA |

It is important to point out again that the quality factor in Table I is weakly affected by body loading, in contrast to conventional high-frequency MRI RF coils. This observation, also mentioned in [16], is a critical advantage of the present electromagnetic stimulator. As to the RF power losses, they are mostly in the coil itself and not in the human body.

B-field measurements have been performed via a calibrated single-axis coil probe located at the coil axis. The measured and theoretical results differ by no more than 10%.

## IV. Device safety

### A. Method of analysis

Safety estimates rely upon the levels of the magnetic field, electric field, and the so-called Specific Absorption Rate (SAR) within the body. The SAR is the heat rate in the body rising due to an imposed electromagnetic field. At the low frequencies considered in this study, the magnetic field is weakly perturbed by the body presence and can therefore be measured in air. However, SAR and electric field measurements cannot be performed easily for human subjects *in vivo*. SAR and device performance estimates are typically derived and accepted today from Computational Electromagnetics (CEM) simulations performed with detailed virtual humans [23]. In this study, we use the multi-tissue CEM phantom VHP-Female v. 5.0 (female/60 year/162 cm/88 kg, obese) [23, 24], derived from the cryosection dataset archived within the Visible Human Project® of the US National Library of Medicine. The phantom includes about 250 individual tissues and is augmented with material property values from the IT’IS database [25]. The average-body conductivity is assigned as 0.25 S/m, which reflects a mixture of muscle and fat.

The primary CEM software used in this study is the accurate commercial FEM solver ANSYS® Electromagnetic Suite 18.2.0 with rigorous adaptive mesh refinement. In addition, and for verification purposes, we employ an in-house boundary element fast multipole method (BEM-FMM) described in [22]. In the latter case, a higher surface resolution can be achieved and the original surface phantom model can be refined and smoothed from approximately 0.5 M triangles to 3.5 M triangles. The human model is placed in the coil at the shoulder landmark, as shown in Fig. 6, so that the top of the shoulder coincides with the ring plane. Other configurations have also been considered. Results obtained with both software packages differ by no more than 2% in the unloaded coil and by no more than 25–50% in the coil loaded with the multi-tissue human body. The latter deviation may be explained by somewhat different surface meshes.

**Fig. 6.**
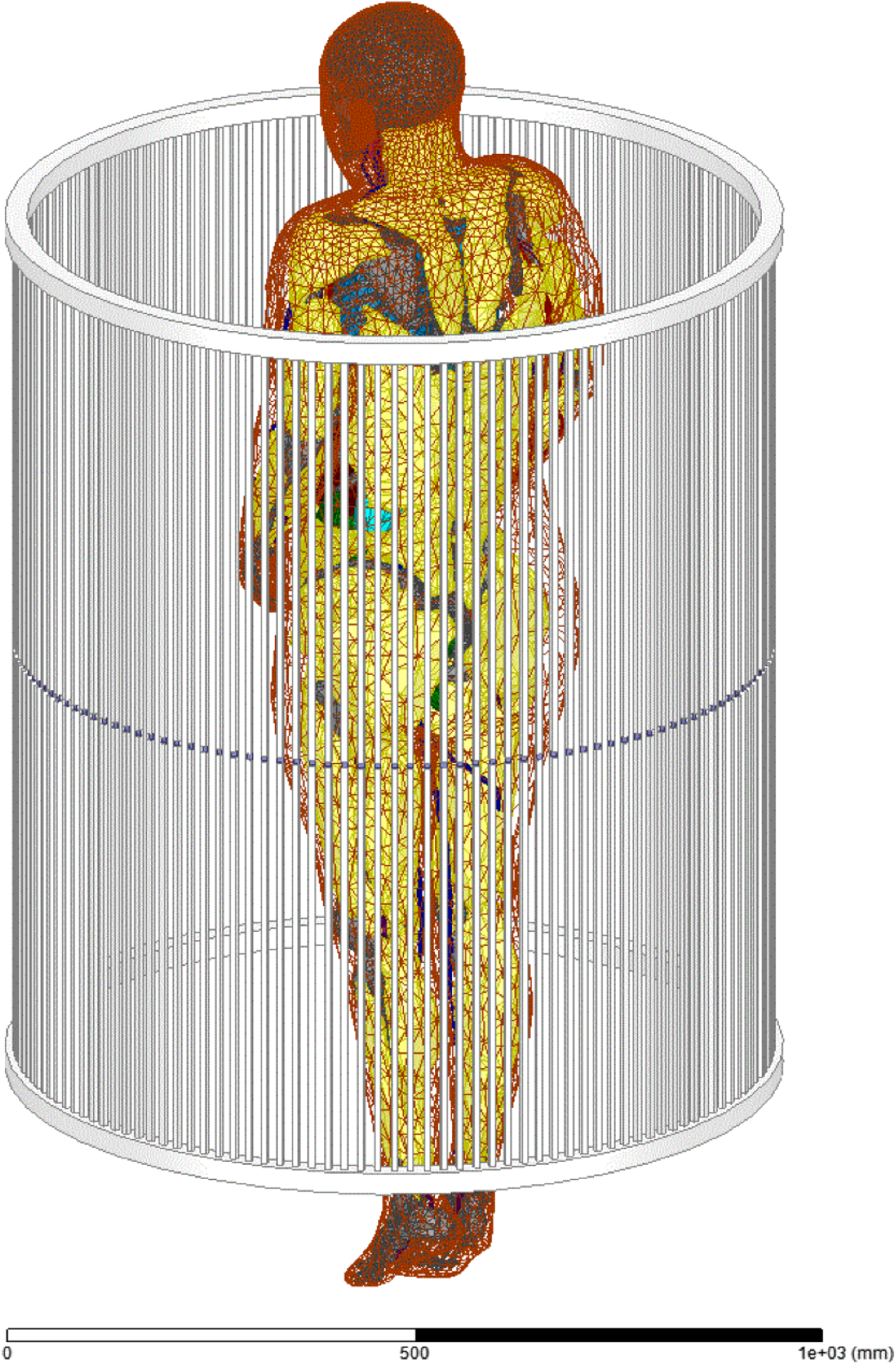
Multi-tissue CEM phantom VHP-Female v. 5.0 within the resonant coil (ANSYS 18.2.0).

Below we report simulations at two power levels: 1.5 kW input power and 3 kW input power. The first power level is the half power level of the amplifier driver; the second power level corresponds to full power. At full power level, the amplitude of the resonant ring current *I*_0_ in Eqs. (4)–(5) reaches 603 A, while the amplitude of the rung current reaches 26 A.

### B. Electric field levels

Guidelines from the International Commission on Non-Ionizing Radiation Protection or ICNIRP, see Table 2 of [26], require the occupational exposure to a time-varying electric field to be limited by a value of 27 V/m RMS at 100 kHz and by a value of 39 V/m RMS at 145 kHz (the so-called basic restrictions [26]). These restrictions are mainly due to the limits on peripheral nerve stimulation.

Fig. 7 shows the simulated RMS levels of the electric field in the body at 100 kHz and at the input power level of 1.5 kW obtained via the BEM-FMM simulations. We observe that, at half power level of the amplifier driver, the fields everywhere in the body do not generally exceed 30 V/m RMS and are thus within the ICNIRP guidelines. As expected, higher field levels are observed closer to the surface; the field gradually decreases toward the center of the body.

**Fig. 7.**
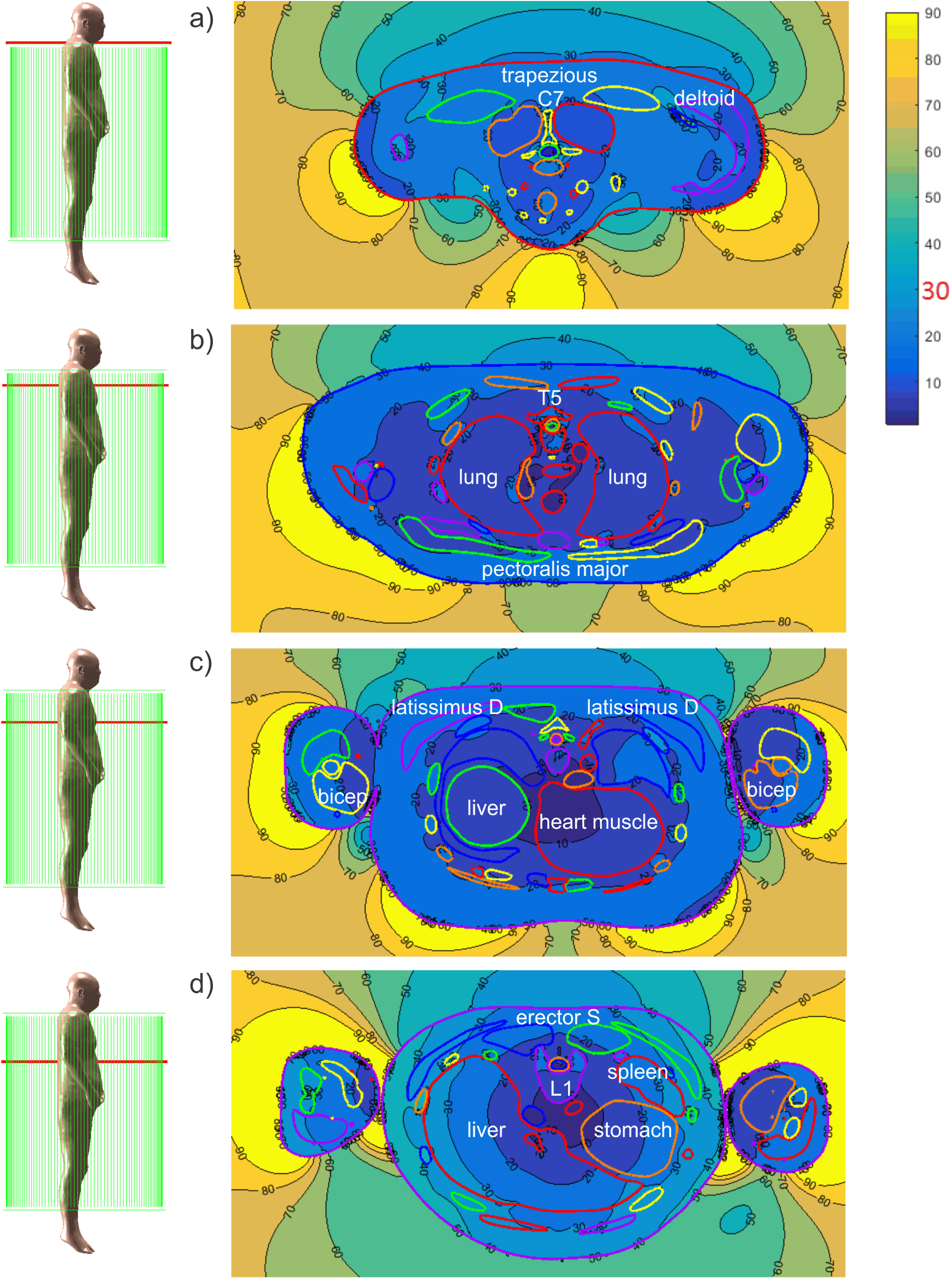
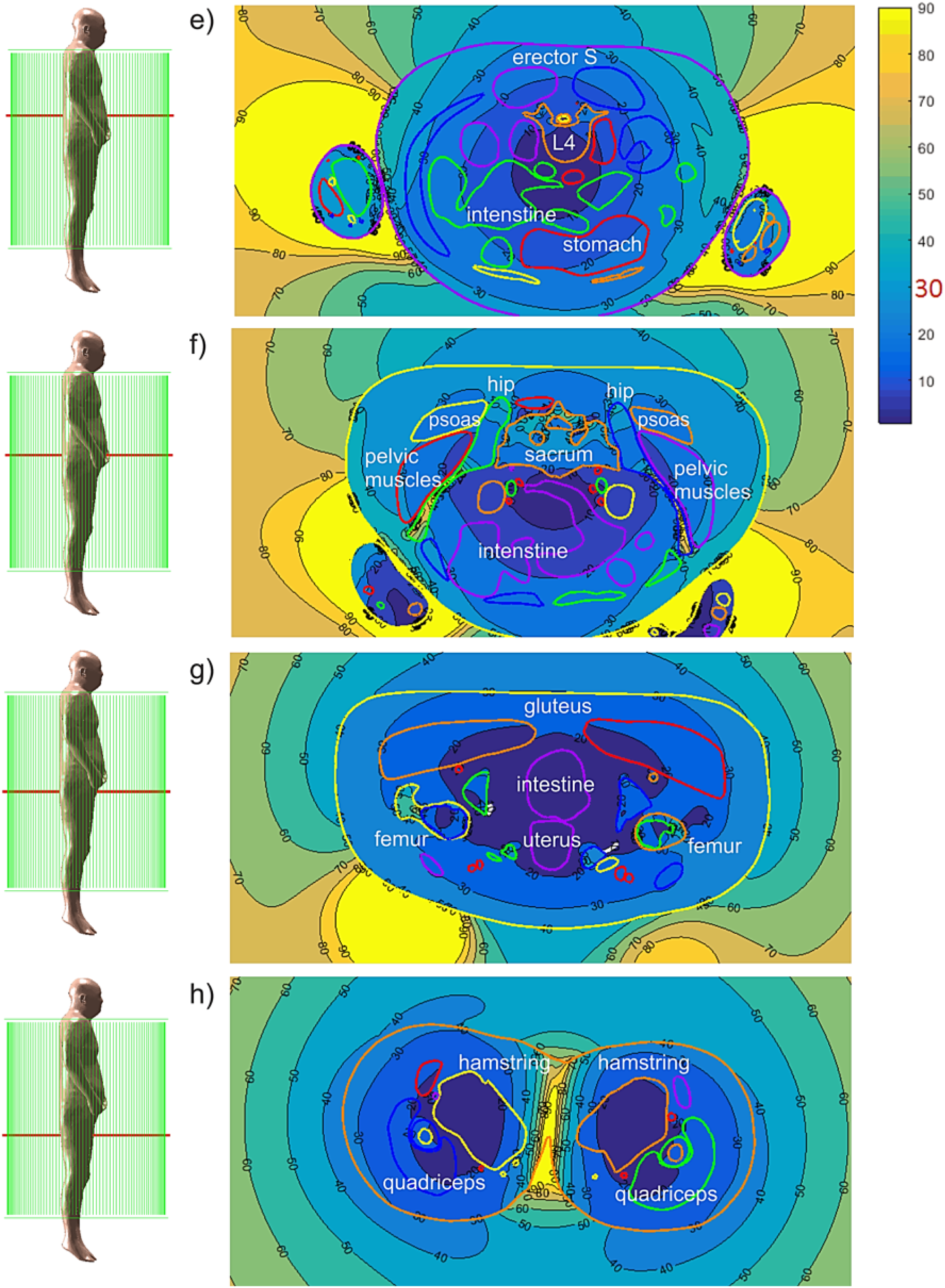
Complex rms magnitude of the electric field (V/m) at 1,500 W input power.

Quantitative estimates of the average electric field for every particular tissue obtained via ANSYS Electromagnetic Suite 18.2.0 are given in Table III. Note the lower electric fields in the intracranial volume. Additionally, we observe higher electric fields in the individual body muscles. It is also interesting to observe that the fields in bone may be quite high, in particular in the femur and pelvic bones.

**TABLE III:** Computed electric field levels (V/m RMS) in every individual tissue at 1.5 kW input power (ANSYS® Electromagnetic Suite 18.2.0).

| Mesh | Tissue | Avg. E Field<br>(V/m RMS) | Mesh | Tissue | Avg. E Field<br>(V/m RMS) |
| --- | --- | --- | --- | --- | --- |
| 1 | Air Internal Maxillary Sinus Left | 7.7 | 39 | Cuneiform Medial right | 0.6 |
| 2 | Air Internal Maxillary Sinus Right | 6.9 | 40 | discC02C03 | 10.3 |
| 3 | Arteries | 10.5 | 41 | discC03C04 | 11.6 |
| 4 | Bladder | 28.0 | 42 | discC04C05 | 14.3 |
| 5 | C01 | 14.5 | 43 | discC05C06 | 16.9 |
| 6 | C02 | 13.6 | 44 | discC06C07 | 19.0 |
| 7 | C03 | 14.8 | 45 | discC07T01 | 20.6 |
| 8 | C04 | 18.5 | 46 | discL01L02 | 13.9 |
| 9 | C05 | 21.7 | 47 | discL02L03 | 11.5 |
| 10 | C06 | 26.1 | 48 | discL03L04 | 8.4 |
| 11 | C07 | 29.6 | 49 | discL04L05 | 4.7 |
| 12 | Calcaneous left | 0.6 | 50 | discL05L06 | 7.1 |
| 13 | Calcaneous right | 1.1 | 51 | discL06S00 | 13.8 |
| 14 | Cartilage1 Left | 18.5 | 52 | discT01T02 | 20.2 |
| 15 | Cartilage1 Right | 19.9 | 53 | discT02T03 | 17.6 |
| 16 | Cartilage2 Left | 19.7 | 54 | discT03T04 | 17.9 |
| 17 | Cartilage2 Right | 20.3 | 55 | discT04T05 | 17.4 |
| 18 | Cartilage3 Left | 21.2 | 56 | discT05T06 | 15.8 |
| 19 | Cartilage3 Right | 20.9 | 57 | discT06T07 | 15.2 |
| 20 | Cartilage4 Left | 22.5 | 58 | discT07T08 | 14.4 |
| 21 | Cartilage4 Right | 21.8 | 59 | discT08T09 | 13.6 |
| 22 | Cartilage5 Left | 24.3 | 60 | discT09T10 | 13.2 |
| 23 | Cartilage5 Right | 22.6 | 61 | discT10T11 | 12.5 |
| 24 | Cartilage6 Left | 36.3 | 62 | discT11T12 | 13.2 |
| 25 | Cartilage6 Right | 35.9 | 63 | discT12L01 | 13.4 |
| 26 | Cerebellum | 1.4 | 64 | Eye Left | 5.0 |
| 27 | Clavicle left | 55.6 | 65 | Eye Right | 5.3 |
| 28 | Clavicle right | 34.9 | 66 | Feet1Phalange left | 0.5 |
| 29 | Coccyx | 42.6 | 67 | Feet1Phalange right | 0.4 |
| 30 | CSF OuterShell | 3.5 | 68 | Feet2Phalange left | 0.4 |
| 31 | CSF Ventricles | 0.4 | 69 | Feet2Phalange right | 0.4 |
| 32 | Cuboid Left | 0.9 | 70 | Feet3Phalange left | 0.3 |
| 33 | Cuboid Right | 0.6 | 71 | Feet3Phalange right | 0.4 |
| 34 | Cuneiform Intermediate left | 1.3 | 72 | Feet4Phalange left | 0.4 |
| 35 | Cuneiform Intermediate right | 0.5 | 73 | Feet4Phalange right | 0.6 |
| 36 | Cuneiform Lateral left | 1.1 | 74 | Feet5Phalange left | 0.4 |
| 37 | Cuneiform Lateral right | 0.4 | 75 | Feet5Phalange right | 0.7 |
| 38 | Cuneiform Medial left | 1.3 | 76 | Femur Bone Marrow Left | 7.1 |
| 77 | Femur Bone Marrow Right | 8.6 | 117 | Humerus right | 23.1 |
| 78 | Femur left | 69.3 | 118 | Intestine | 20.6 |
| 79 | Femur right | 83.7 | 119 | Jaw lower | 10.0 |
| 80 | Fibula left | 5.9 | 120 | Kidney left | 29.3 |
| 81 | Fibula right | 5.4 | 121 | Kidney right | 27.2 |
| 82 | Grey Matter Spinal Cord | 1.5 | 122 | L01 | 27.9 |
| 83 | Hands1 1Phalange left | 10.2 | 123 | L02 | 25.3 |
| 84 | Hands1 1Phalange right | 9.5 | 124 | L03 | 22.1 |
| 85 | Hands1 2Phalange left | 9.5 | 125 | L04 | 19.0 |
| 86 | Hands1 2Phalange right | 12.1 | 126 | L05 | 14.4 |
| 87 | Hands1 3Phalange left | 12.2 | 127 | L06 | 17.9 |
| 88 | Hands1 3Phalange right | 13.4 | 128 | Liver | 29.8 |
| 89 | Hands2 1Phalange left | 7.6 | 129 | Lungs | 19.4 |
| 90 | Hands2 1Phalange right | 7.1 | 130 | Median Nerve left | 11.6 |
| 91 | Hands2 2Phalange left | 8.1 | 131 | Median Nerve right | 13.0 |
| 92 | Hands2 2Phalange right | 8.5 | 132 | Muscle Bicep left | 11.9 |
| 93 | Hands2 3Phalange left | 7.1 | 133 | Muscle Bicep right | 12.9 |
| 94 | Hands2 3Phalange right | 9.5 | 134 | Muscle Calf left | 5.0 |
| 95 | Hands3 1Phalange left | 6.4 | 135 | Muscle Calf right | 5.2 |
| 96 | Hands3 1Phalange right | 6.2 | 136 | Muscle Deltoid left | 18.7 |
| 97 | Hands3 2Phalange left | 8.1 | 137 | Muscle Deltoid right | 19.3 |
| 98 | Hands3 2Phalange right | 8.6 | 138 | Muscle Erector spinae left | 26.8 |
| 99 | Hands3 3Phalange left | 9.3 | 139 | Muscle Erector spinae right | 26.9 |
| 100 | Hands3 3Phalange right | 11.0 | 140 | Muscle Forearm Extensors left | 6.9 |
| 101 | Hands4 1Phalange left | 7.2 | 141 | Muscle Forearm Extensors right | 8.6 |
| 102 | Hands4 1Phalange right | 6.9 | 142 | Muscle Forearm Flexors left | 6.9 |
| 103 | Hands4 2Phalange left | 10.4 | 143 | Muscle Forearm Flexors right | 7.2 |
| 104 | Hands4 2Phalange right | 10.0 | 144 | Muscle Gluteus left | 27.9 |
| 105 | Hands4 3Phalange left | 11.1 | 145 | Muscle Gluteus right | 27.2 |
| 106 | Hands4 3Phalange right | 10.7 | 146 | Muscle Hamstring left | 18.9 |
| 107 | Hands5 1Phalange left | 9.0 | 147 | Muscle Hamstring right | 19.1 |
| 108 | Hands5 1Phalange right | 10.3 | 148 | Muscle Latissimus Dorsi left | 36.6 |
| 109 | Hands5 2Phalange left | 11.0 | 149 | Muscle Latissimus Dorsi right | 38.5 |
| 110 | Hands5 2Phalange right | 12.6 | 150 | Muscle Neck Combined left | 13.6 |
| 111 | Hands5 3Phalange left | 10.3 | 151 | Muscle Neck Combined right | 13.4 |
| 112 | Hands5 3Phalange right | 12.1 | 152 | Muscle Obliques left | 39.5 |
| 113 | Heart Muscle | 14.2 | 153 | Muscle Obliques right | 40.1 |
| 114 | Hip left | 60.0 | 154 | Muscle Pectoralis major left | 21.7 |
| 115 | Hip right | 61.4 | 155 | Muscle Pectoralis major right | 20.9 |
| 116 | Humerus left | 20.7 | 156 | Muscle Pectoralis minor left | 19.2 |
| 157 | Muscle Pectoralis minor right | 18.6 | 194 | Ribs left8 | 47.3 |
| 158 | Muscle Pelvic Combined left | 25.7 | 195 | Ribs left9 | 46.1 |
| 159 | Muscle Pelvic Combined right | 25.0 | 196 | Ribs left10 | 48.5 |
| 160 | Muscle Psoas left | 13.9 | 197 | Ribs left11 | 51.7 |
| 161 | Muscle Psoas right | 13.9 | 198 | Ribs left12 | 39.0 |
| 162 | Muscle Quadriceps left | 20.2 | 199 | Ribs right1 | 29.6 |
| 163 | Muscle Quadriceps right | 19.9 | 200 | Ribs right2 | 26.2 |
| 164 | Muscle Rectus Abdominis left bottom | 32.3 | 201 | Ribs right3 | 25.2 |
| 165 | Muscle Rectus Abdominis left middle | 34.9 | 202 | Ribs right4 | 26.1 |
| 166 | Muscle Rectus Abdominis left top | 39.1 | 203 | Ribs right5 | 27.2 |
| 167 | Muscle Rectus Abdominis right bottom | 32.5 | 204 | Ribs right6 | 29.9 |
| 168 | Muscle Rectus Abdominis right middle | 35.4 | 205 | Ribs right7 | 35.6 |
| 169 | Muscle Rectus Abdominis right top | 38.1 | 206 | Ribs right8 | 43.2 |
| 170 | Muscle Sartorius left | 18.6 | 207 | Ribs right9 | 53.9 |
| 171 | Muscle Sartorius right | 17.5 | 208 | Ribs right10 | 58.9 |
| 172 | Muscle Tibialis Anterior left | 6.2 | 209 | Ribs right11 | 56.9 |
| 173 | Muscle Tibialis Anterior right | 5.8 | 210 | Ribs right12 | 40.9 |
| 174 | Muscle Trapezius left | 23.6 | 211 | Sacrum | 45.7 |
| 175 | Muscle Trapezius right | 24.0 | 212 | Scapula left | 38.2 |
| 176 | Muscle Tricep left | 12.0 | 213 | Scapula right | 38.8 |
| 177 | Muscle Tricep right | 14.0 | 214 | Skin Shell | 27.8 |
| 178 | Navicular left | 1.8 | 215 | Skull | 22.8 |
| 179 | Navicular right | 0.7 | 216 | Sphenoid | 8.9 |
| 180 | Patella left | 24.3 | 217 | Spleen | 33.9 |
| 181 | Patella right | 22.6 | 218 | Sternum | 25.2 |
| 182 | Peripheral Nerve left | 17.1 | 219 | Stomach | 22.6 |
| 183 | Peripheral Nerve Right | 14.1 | 220 | T01 | 28.4 |
| 184 | Pubic Symphysis | 32.1 | 221 | T02 | 27.2 |
| 185 | Radial Nerve left | 14.6 | 222 | T03 | 27.2 |
| 186 | Radial Nerve right | 12.4 | 223 | T04 | 26.9 |
| 187 | Ribs left1 | 26.4 | 224 | T05 | 25.7 |
| 188 | Ribs left2 | 30.1 | 225 | T06 | 25.5 |
| 189 | Ribs left3 | 26.4 | 226 | T07 | 26.2 |
| 190 | Ribs left4 | 26.7 | 227 | T08 | 26.4 |
| 191 | Ribs left5 | 28.3 | 228 | T09 | 26.9 |
| 192 | Ribs left6 | 31.5 | 229 | T10 | 26.1 |
| 193 | Ribs left7 | 37.3 | 230 | T11 | 26.4 |
| 231 | T12 | 27.5 | 240 | Trabecular upper right | 0.9 |
| 232 | Talus left | 1.3 | 241 | Trachea Sinus | 12.4 |
| 233 | Talus right | 0.6 | 242 | Ulna Radius left | 8.1 |
| 234 | Tibia left | 8.3 | 243 | Ulna Radius right | 7.8 |
| 235 | Tibia right | 7.9 | 244 | Uterus | 17.3 |
| 236 | Tongue | 5.2 | 245 | Veins lower | 12.5 |
| 237 | Trabecular lower left | 0.5 | 246 | Veins upper | 12.4 |
| 238 | Trabecular lower right | 0.8 | 247 | White Matter | 1.0 |
| 239 | Trabecular upper left | 0.7 |  |  |  |

However, the computed local electric fields may considerably exceed the values reported in Table III, in particular by 1.5–6 times. These peak values are less accurate. One potential source of the numerical error is insufficient resolution of lengthy and time-consuming full-body computations very close to the interfaces where higher fields are usually observed.

### C. SAR levels

The body-averaged or the whole-body (global-body) SAR_body_ is given by averaging the local SAR over the entire body volume. In terms of the complex field phasor **E**(**r**), one has

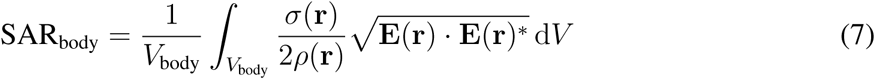

Here, *σ*(**r**) is the local tissue conductivity and *ρ*(**r**) is the local mass density. At full power of 3 kW and positioned at the shoulder landmark, the global-body SAR computed via ANSYS Electromagnetic Suite 18.2.0 is 0.25 W/kg. Thus, the total power dissipation in the body does not exceed 30 W, i.e., 1% of the total power. The same percentage ratio is valid at half input power.

The second critical estimate is SAR_1 g_, which is given by averaging over a contiguous volume with the weight of 1 g,

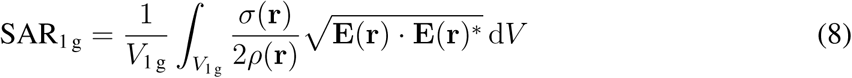

The maximum value of SAR_1 g_, in the body computed via ANSYS Electromagnetic Suite 18.2.0 at the full power of 3 kW and located at the shoulder landmark is 4.55 W/kg.

Although this last value might appear to be relatively high, it is still within the corresponding SAR limits in MRI machines [27, 28]. In particular, the major applicable MRI safety standard [28] issued by the International Electrotechnical Commission (IEC) and also accepted by the U.S. Food and Drug Administration in the normal mode (mode of operation that causes no physiological stress to patients) limits global-body SAR to 2 W/kg, global-head SAR to 3.2 W/kg, local head and torso SAR to 10 W/kg, and local extremity SAR to 20 W/kg [27]. The global SAR limits are intended to ensure a body core temperature of 39°C or less [27, 28].

## V. Discussion

We envision several potential application scenarios of the present electromagnetic stimulation device, though others may certainly be feasible. First, it could be tested for chronic pain treatment as suggested in the introduction. For example, one potential target is the spinal cord. Epidural spinal cord stimulation (SCS) has been used for pain management in the past several decades [29]. However, SCS involves surgical implantation of a pulse generator device that delivers weak currents to nerve fibers of the spinal cord. A noninvasive alternative is transcutaneous spinal direct current stimulation (tcDCS) [30], which delivers an average electric field of 0.15 V/m in the spinal cord between the electrodes. In comparison, our coil can induce an electric field in the spinal cord grey matter of approximately an order of magnitude higher compared to direct current stimulation.

Another potential application for our coil device is treatment of psychiatric disorders such as major depressive disorder. Repetitive transcranial magnetic stimulation (rTMS) was first shown to be efficacious for the treatment of depression in the mid-1990s [31] and subsequently cleared by the U.S. FDA in 2008 for treatment-resistant depression (TRD). Clinical rTMS uses a focal figure-of-8 coil to deliver electric field pulses at the dorsal lateral prefrontal cortex at an intensity sufficient to induce action potentials in the underlying neurons. Several strategies using nonfocal (whole brain) subthreshold stimulation are currently being explored for the treatment of TRD. For example, it was reported that low-field (*<* 1 V/m) stimulation delivered using a MRI-gradient type coil had a rapid mood-elevating effect in animals and bipolar patients [32–35], and has been shown to affect brain glucose metabolism [36]. In addition, a system has been proposed, using a series of rotating permanent magnets to induce a small current in the brain in order to entrain neural oscillations, enhance cortical plasticity, normalize cerebral blood flow, and altogether ameliorate depressive symptoms [37]. An embodiment of our coil can be made smaller for efficient transcranial brain stimulation.

## VI. Conclusions

In this technical study we described a whole-body non-contact electromagnetic stimulation device based on the concept of a familiar MRI RF resonating coil, but at a much lower resonant frequency (100–150 kHz), with a field modulation option (0.5–100 Hz) and with an input power level of up to 3 kW. Its unique features include a high electric field level within the subject’s biological tissue due to the resonant effect but at a low power dissipation, or SAR level, in the body itself.

Due to a large resonator volume and its non-contact nature, the subject may be conveniently located anywhere within the resonating coil over a prolonged period of time at moderate and safe electric field levels. The electric field effect does not depend on a particular body position within the resonator. The field penetration is deep everywhere in the body including the extremities; muscles, bones, and peripheral tissues are mostly affected. Over a shorter period of time, the electric field levels could be increased to relatively large values with an amplitude of about 1 V/cm.

We envision treatment of chronic pain, and particularly neuropathic pain, as the primary potential clinical application for the device. The device enables whole-body coverage, which could be useful in the treatment of widespread pain conditions, such as painful polyneuropathy or fibromyalgia. In addition, a deeper tissue penetration can be achieved without causing side-effects caused by high current density in the skin associated with the traditional contact electrodes. Another potential application might include for example facilitation of chronic skin wound healing [38]. Medium to high electric field levels approaching or even exceeding 30 V/m RMS could likely be applied. It should be noted that these potential clinical applications are speculative and warrant empirical testing in the future.

Considerable attention has been paid to device safety including both the AC power safety and human exposure to electromagnetic fields. In the former case, we have used inductive coupling, which assures that there is no direct current path from the AC power outlet to the coil. This design enhances overall device safety at any power level, including high-power operation. As with more traditional MRI devices, no large metal objects should be located in the immediate vicinity of the coil.

Human exposure to the electromagnetic field within the coil has been evaluated by performing extensive modeling with two independent numerical methods and with an anatomically realistic multi-tissue human phantom. We have shown that the SAR levels within the body correspond to the safety standards of the International Electrotechnical Commission when the input power level of the amplifier driver does not exceed 3 kW. We have also shown that the electric field levels generally comply with the safety standards of the International Commission on Non-Ionizing Radiation Protection when the input power level of the amplifier driver does not exceed 1.5 kW.

## Acknowledgments

The authors are thankful to Dr. James O’Rourke, Dr. John McNeill, Ms. Leah Morales, and Mr. Brandon Weyant (all from Worcester Polytechnic Institute), MD Irina V. Zhdanova (ClockCoach), and Dr. Aapo Nummenmaa (Massachusetts General Hospital) for useful discussions. Dr. Deng is supported by the Intramural Research Program of the National Institute of Mental Health, NIH.

